# Local Connectome Phenotypes Predict Social, Health, and Cognitive Factors

**DOI:** 10.1101/122945

**Authors:** Michael A. Powell, Javier O. Garcia, Fang-Cheng Yeh, Jean M. Vettel, Timothy Verstynen

## Abstract

The unique architecture of the human connectome is defined initially by genetics and subsequently sculpted over time with experience. Thus, similarities in predisposition and experience that lead to similarities in social, biological, and cognitive attributes should also be reflected in the local architecture of white matter fascicles. Here we employ a method known as local connectome fingerprinting that uses diffusion MRI to measure the fiber-wise characteristics of macroscopic white matter pathways throughout the brain. This fingerprinting approach was applied to a large sample (N=841) of subjects from the Human Connectome Project, revealing a reliable degree of between-subject correlation in the local connectome fingerprints, with a relatively complex, low-dimensional substructure. Using a cross-validated, high-dimensional regression analysis approach, we derived local connectome phenotype (LCP) maps that could reliably predict a subset of subject attributes measured, including demographic, health and cognitive measures. These LCP maps were highly specific to the attribute being predicted but also sensitive to correlations between attributes. Collectively, these results indicate that the local architecture of white matter fascicles reflects a meaningful portion of the variability shared between subjects along several dimensions.

**Author Summary:** The local connectome is the pattern of fiber systems (i.e., number of fibers, orientation, and size) within a voxel and reflects the proximal characteristics of white matter fascicles distributed throughout the brain. Here we show how variability in the local connectome is correlated in a principled way across individuals. This inter-subject correlation is reliable enough that unique phenotype maps can be learned to predict between-subject variability in a range of social, health, and cognitive attributes. This work shows, for the first time, how shared variability across individuals is reflected in the local connectome.

## Introduction

The unique pattern of connections among the billions of neurons in the brain is termed the connectome (Sporns, Tononi, & Kotter, 2005), and this pattern encapsulates a fundamental constraint on neural computation and cognition (Gu et al., 2015; Thivierge & Marcus, 2007). This connective architecture is initially structured by genetics and then sculpted by experience over time (Kochunov, Fu, et al., 2016; Kochunov, Thompson, et al., 2016; Yeh, Vettel, et al., 2016). Recent advancements in neuroimaging techniques, particularly diffusion MRI (dMRI), have opened the door to mapping the macroscopic-level properties of the structural connectome in vivo (Le Bihan & Johansen-Berg, 2012). As a result, a growing body of research has focused on quantifying how variability in structural connectivity associates with individual differences in functional properties of brain networks (Muldoon et al., 2016; Passingham, Stephan, & Kötter, 2002), as well as associating with differences in social (Gianaros, Marsland, Sheu, Erickson, & Verstynen, 2013; Molesworth, Sheu, Cohen, Gianaros, & Verstynen, 2015), biological (Arfanakis et al., 2013; Miralbell et al., 2012; Verstynen et al., 2013), and cognitive (Muraskin et al., 2016; Verstynen, 2014; Ystad et al., 2011) attributes.

DMRI works by measuring the microscopic diffusion pattern of water trapped in cellular tissues, allowing for a full characterization of white matter pathways, such as axonal fiber direction and integrity (for review see Jbabdi, Sotiropoulos, Haber, Van Essen, & Behrens, 2015; Le Bihan & Johansen-Berg, 2012). Previous studies have used dMRI to map the global properties of the macroscopic connectome by determining end-to-end connectivity between brain regions (Hagmann et al., 2010; Hagmann et al., 2008, Hagmann 2010; Sporns, 2014). The resulting connectivity estimates can then be summarized, often using graph theoretic techniques that are then associated with variability across individuals (Bullmore & Sporns, 2009; Rubinov & Sporns, 2010). While dMRI acquisition and reconstruction approaches have improved substantially in recent years (Fan et al., 2016; Van Essen et al., 2012), the reliability and validity of many popular fiber tractography algorithms have come into question (Daducci, Dal Palú, Descoteaux, & Thiran, 2016; Reveley et al., 2015; Thomas et al., 2014). As a result, the reliability of subsequent inter-region connectivity estimates may be negatively impacted.

Instead of mapping end-to-end connectivity between regions, we recently introduced the concept of the local connectome as an alternative measure of structural connectivity that does not rely on fiber tracking (Yeh, Badre, & Verstynen, 2016). The local connectome is defined as the pattern of fiber systems (i.e., number of fibers, orientation, and size) within a voxel, as well as immediate connectivity between adjacent voxels, and can be quantified by measuring the fiber-wise density of microscopic water diffusion within a voxel. This voxel-wise measure shares many similarities with the concept of a “fixel” proposed by others (Raffelt et al., 2015). The complete collection of these multi-fiber diffusion density measurements within all white matter voxels, termed the local connectome fingerprint, provides a high-dimensional feature vector that can describe the unique configuration of the structural connectome (Yeh, Vettel, et al., 2016). In this way, the local connectome fingerprint provides a diffusion-informed measure along the fascicles that supports inter-regional communication, rather than determining the start and end positions of a particular fiber bundle.

We recently showed that the local connectome fingerprint is highly specific to an individual, affording near-perfect accuracy on within-versus-between subject classification tests among hundreds of participants (Yeh, Badre, et al., 2016). Importantly, this demonstrated that a large portion of an individual’s local connectome is driven by experience. Whole-fingerprint distance tests revealed only a 12.51% similarity between monozygotic twins, relative to almost no similarity between genetically unrelated individuals. In addition, within-subject uniqueness showed substantial plasticity, changing at a rate of approximately 12.79% every 100 days (Yeh, Vettel, et al., 2016). Thus, the unique architecture of the local connectome appears to be initially defined by genetics and then subsequently sculpted over time with experience.

The plasticity of the local white matter architecture suggests that it is important to consider how whole-fingerprint uniqueness may mask more subtle similarities arising from common experiences. If experience, including common social or environmental factors, is a major force impacting the structural connectome, then common experiences between individuals may also lead to increased similarity in their local connectomes. In addition, since the white matter is a fundamental constraint on cognition, similarities in local connectomes are expected to associate with similarities in cognitive function. Thus, we hypothesized that shared variability in certain social, biological, or cognitive attributes can be predicted from the local connectome fingerprints.

To test this, we reconstructed multi-shell dMRI data from the Human Connectome Project (HCP) to produce individual local connectome fingerprints from 841 subjects. A set of 32 subject-level attributes was used for predictive modeling, including many social, biological, and cognitive factors. A model between each fiber in the local connectome fingerprint and a target attribute was learned using a cross-validated, sparse version of principal component regression. The predictive utility of each attribute map, termed a local connectome phenotype (LCP), was evaluated by predicting a given attribute using cross validation. Our results show that specific characteristics of the local connectome are sensitive to shared variability across individuals, as well as being highly reliable within an individual (Yeh, Vettel, et al., 2016), confirming its utility for understanding how network organization reflects genetic and experiential factors.

## Materials and Methods

### Participants

We used publicly available dMRI data from the S900 (2015) release of the Human Connectome Project (HCP; (Van Essen et al., 2013)), acquired by Washington University in St. Louis and the University of Minnesota. Out of the 900 participants released, 841 participants (370 male, ages 22-37, mean age 28.76) had viable dMRI datasets. Our analysis was restricted to this subsample. All data collection procedures were approved by the institutional review boards at Washington University in St. Louis and the University of Minnesota. The post hoc data analysis was approved as exempt by the institutional review board at Carnegie Mellon University, in accordance with 45 CFR 46.101(b)(4) (IRB Protocol Number: HS14-139).

### Diffusion MRI Acquisition

The dMRI data were acquired on a Siemens 3T Skyra scanner using a 2D spin-echo single-shot multiband EPI sequence with a multiband factor of 3 and monopolar gradient pulse. The spatial resolution was 1.25 mm isotropic (TR = 5500 ms, TE = 89.50 ms). The b-values were 1000, 2000, and 3000 s/mm^2^. The total number of diffusion sampling directions was 90 for each of the three shells in addition to 6 b0 images. The total scanning time was approximately 55 minutes.

### Local Connectome Fingerprint Reconstruction

An outline of the pipeline for generating local connectome fingerprints is shown in the top panel of Figure 1. The dMRI data for each subject was reconstructed in a common stereotaxic space using q-space diffeomorphic reconstruction (QSDR) (F. C. Yeh & Tseng, 2011), a nonlinear registration approach that directly reconstructs water diffusion density patterns into a common stereotaxic space at 1-mm^3^ resolution.

Using the HCP dataset, we derived an atlas of axonal direction in each voxel (publicly available at http://dsi-studio.labsolver.org). A spin distribution function (SDF) sampling framework was used to provide a consistent set of directions 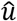 to sample the magnitude of SDFs along axonal directions in the cerebral white matter. Since each voxel may have more than one fiber direction, multiple measurements were extracted from the SDF for voxels that contained crossing fibers, while a single measurement was extracted for voxels with fibers in a single direction. The appropriate number of density measurements from each voxel was sampled by the left-posterior-superior voxel order and compiled into a sequence of scalar values. Gray matter was excluded using the ICBM-152 white matter mask (MacConnell Brain Imaging Centre, McGill University, Canada). The cerebellum was also excluded due to different slice coverage in cerebellum across participants. Since the density measurement has arbitrary units, the local connectome fingerprint was scaled to make the variance equal to 1 (Yeh, Vettel, et al., 2016). The resulting local connectome fingerprint is thus a one-dimensional vector where each entry represents the density estimate of restricted water diffusion in a specific direction along an average fiber. The magnitude of this value reflects the average signal across a large number of coherently oriented axons, as well as support tissue like myelin and other glia.

The local connectome fingerprint construction was conducted using DSI Studio (http://dsi-studio.labsolver.org), an open-source diffusion MRI analysis tool for connectome analysis. The source code, documentation, and local connectome fingerprint data are publicly available on the same website.

**Figure 1.**
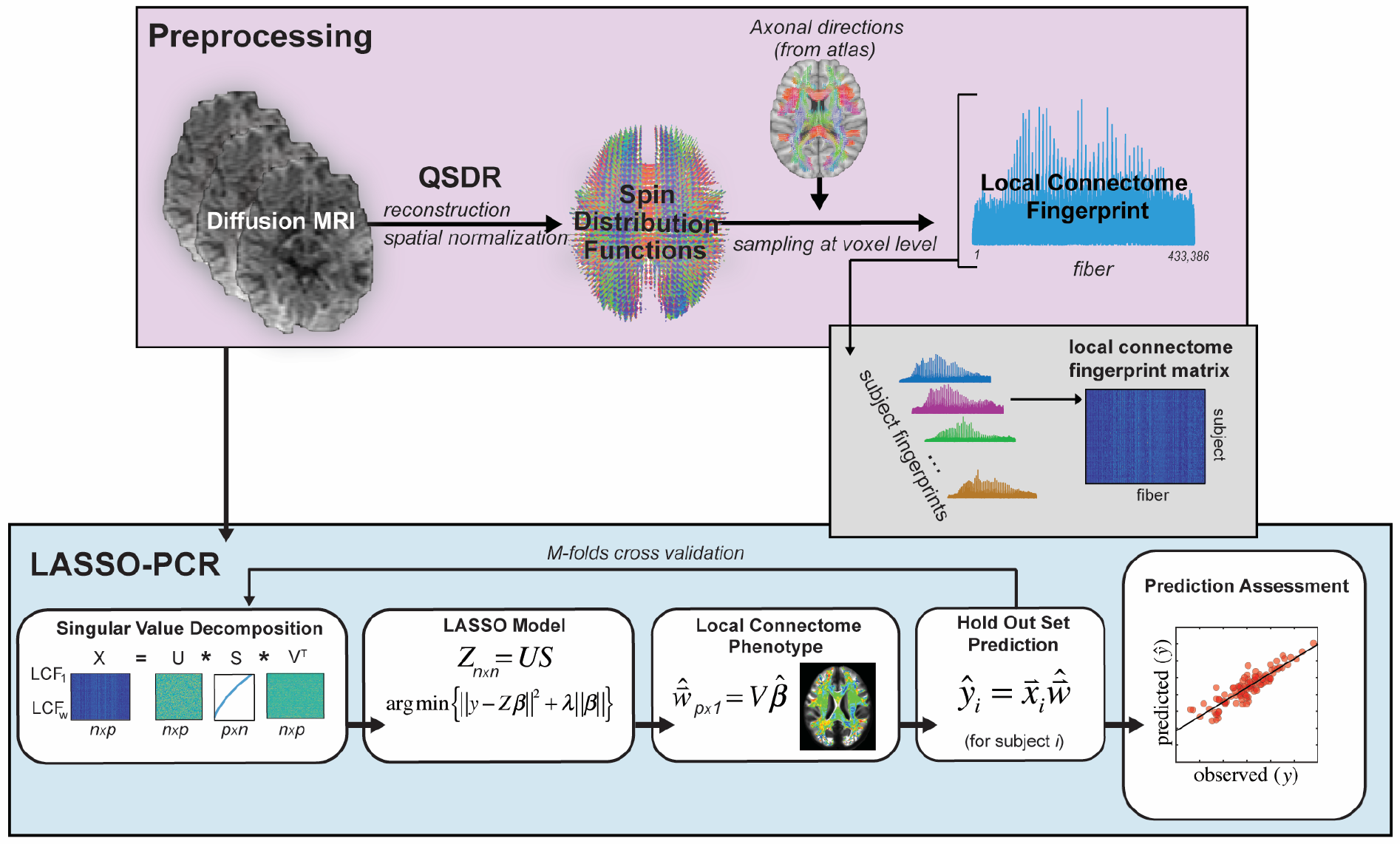
Data analysis pipeline. dMRI from the HCP dataset were preprocessed consistent with previous research investigating the local connectome fingerprint (top panel) and included registration via QSDR and estimation of SDF using an axonal directional atlas derived from the HCP dataset. Once fingerprints were estimated for each individual, the pipeline for analysis of the continuous response variables consisted of four major steps: 1) a PCA-based dimensionality reduction, 2) a LASSO model based on the lower-dimensional components of the local connectome fingerprint, 3) local connectome phenotype estimation from projection of the contributing components of the LASSO model, and 4) prediction on the held-out dataset. A similar pipeline was used for categorical response variables with the exception that a logistic lasso model was used in the LASSO-PCR step and prediction accuracy was assessed as percent correct rather than as a predicted vs. observed correlation.

### Response Variables

A total of 32 response variables across social, health, and cognitive factors were selected from the public and restricted data sets released as part of the HCP. Each variable is summarized in Table 1, but additional details can be found in the HCP Data Dictionary (https://wiki.humanconnectome.org/display/PublicData/HCP+Data+Dictionary+Public-+500+Subject+Release). Table 1 provides a description of relevant distributional parameters of all of the continuous variables tested. Descriptions of distributional properties of categorical variables are provided in the descriptions below. Supplementary Table 1 shows the correlation between all continuous variables tested.

Demographic and social factors included age (years), gender (56% female, 44% male), race (82% white and 18% black in a reduced subset of the total population), ethnicity (91.4% Hispanic, 8.6% non-Hispanic), handedness, income (from the Semi-Structured Assessment for the Genetics of Alcoholism (SSAGA) scale), education (SSAGA), and relationship status (SSAGA, 44.3% in a "Married or Live-in Relationship" and 55.7% not in such a relationship).

Health factors included body mass index, mean hematocrit, blood pressure (diastolic and systolic), hemoglobin A1c, and sleep quality (Pittsburgh Sleep Quality Index).

Cognitive measures included 11 tests that sampled a broad spectrum of domains: (1) the NIH Picture Sequence Memory Test assessed episodic memory performance, (2) NIH Dimensional Change Card Sort tested executive function and cognitive flexibility, (3) NIH Flanker Inhibitory Control and Attention Test evaluated executive function and inhibition control, (4) Penn Progressive Matrices examined fluid intelligence and was measured using three performance metrics (number of correct responses, total skipped items, and median reaction time for correct responses), (5) NIH Oral Reading Recognition Test assessed language and reading performance, (6) NIH Picture Vocabulary Test examined language skills indexed by vocabulary comprehension, (7) NIH Pattern Comparison Processing Speed Test evaluated processing speed, (8) Delay Discounting tested self-regulation and impulsivity control using two different financial incentives (Area Under the Curve (AUC) for discounting of $200, AUC for discounting of $40,000), (9) Variable Short Penn Line Orientation assessed spatial orientation performance and was measured using three metrics (total number correct, median reaction time divided by expected number of clicks for correct, total positions off for all trials), (10) Penn Word Memory Test evaluated verbal episodic memory using two performance metrics (total number of correct responses, median reaction time for correct responses), and (11) the NIH List Sorting Task tested working memory performance.

**Table 1.**
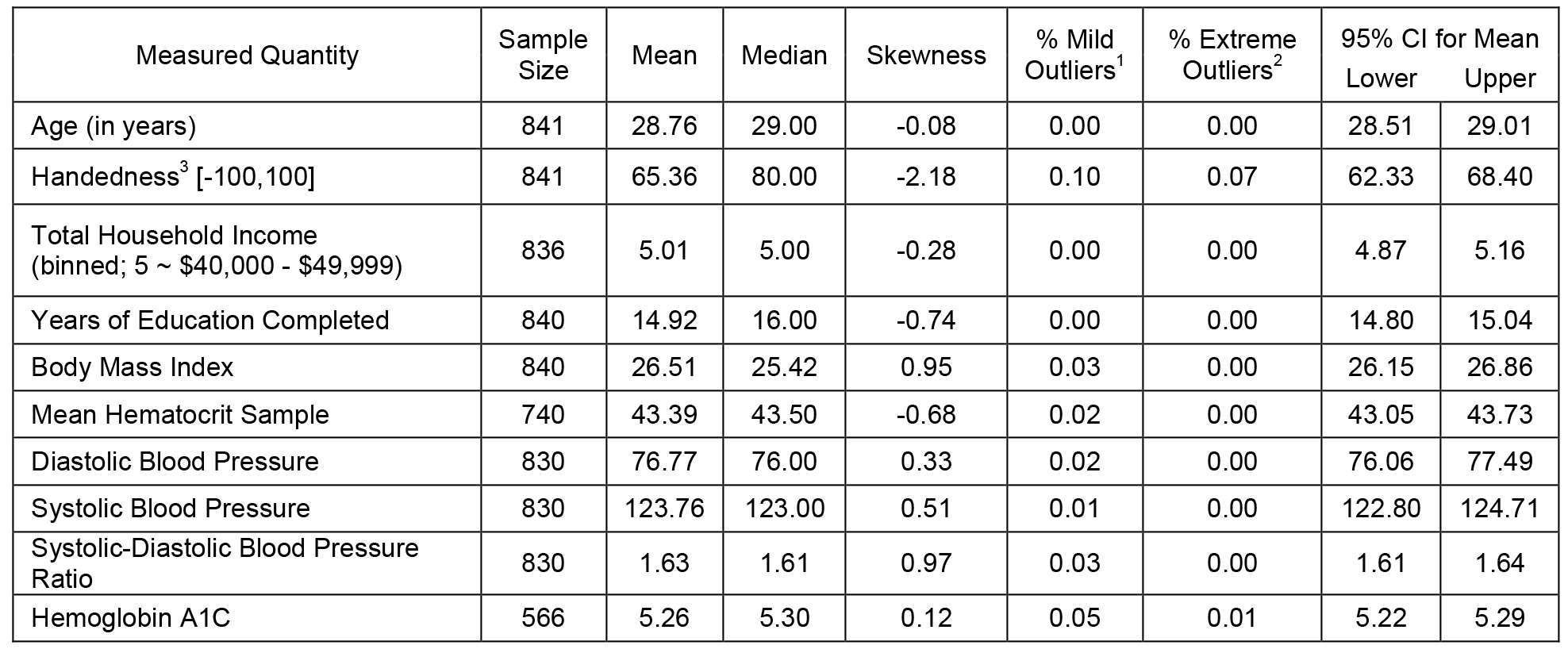
Summary statistics for 28 continuous HCP attributes used in the modeling analysis.

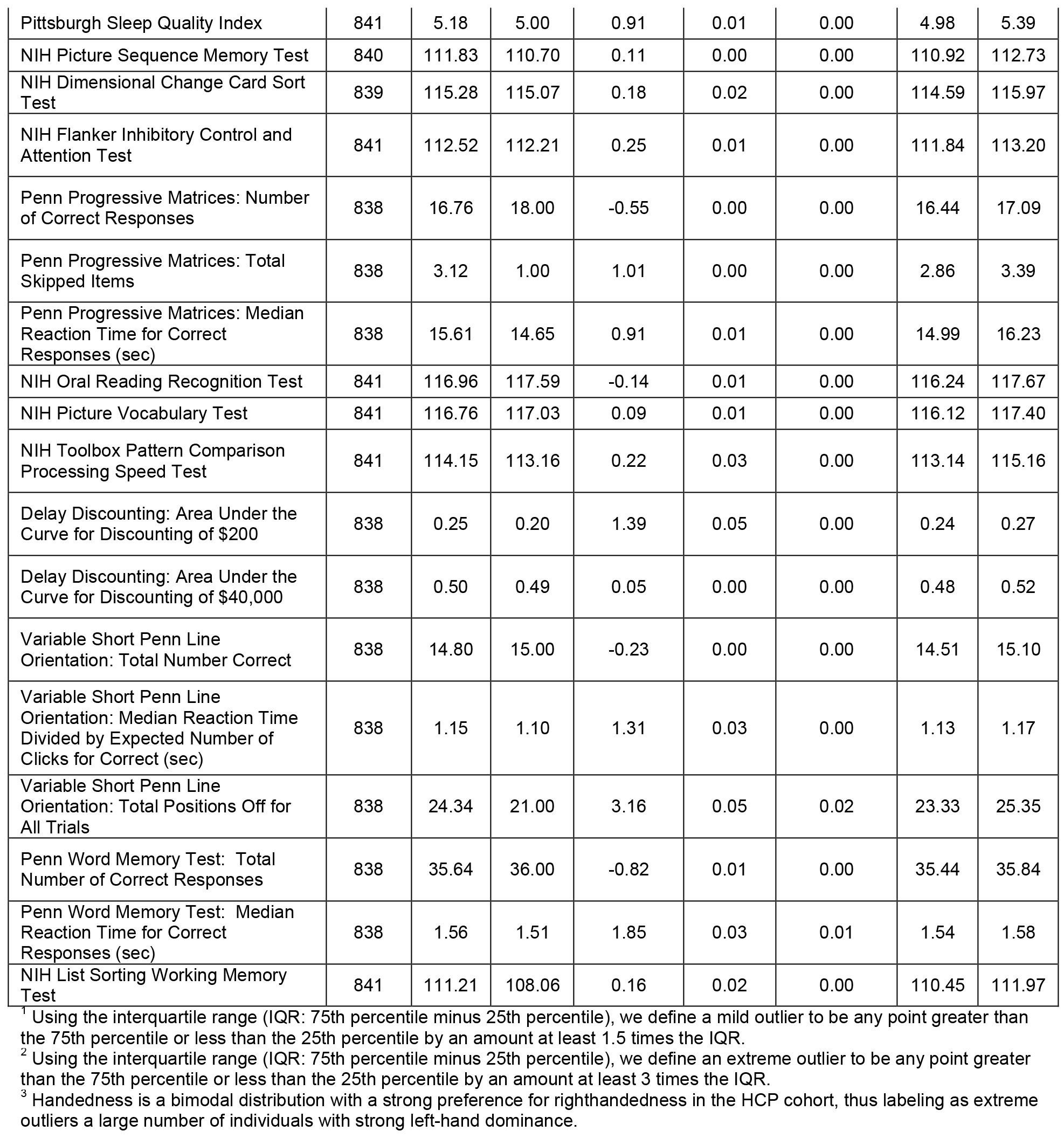

### LASSO Principal Components Regression (LASSO-PCR)

The primary goal of our analysis pipeline was to identify specific patterns of variability in the local connectome that reliably predict individual differences in a specific attribute. These unique patterns would reflect a local connectome phenotype for that attribute. The LASSO-PCR pipeline used to generate local connectome phenotype (LCP) maps is illustrated in the lower panel of Figure 1. This process relied on a 5-fold cross-validation scheme in which a unique 20% of the participants were assigned to each of five subsamples. For each cross-validation fold, we trained models using 80% of the participants in order to make predictions on the held-out 20% of participants. The large number of HCP participants and the infrequent occurrence of outliers in the continuous response variables (see Table 1) justified random fold assignments with little concern about a higher density of outliers existing in any one fold. The random assignment of subjects to folds could pose issues for any infrequent categories in the binary response variables, but the removal of insufficiently represented categories and a verification of near-even class distributions in each fold alleviated these concerns. The analysis pipeline consisted of four major steps.

<u>Step 1: Dimensionality Reduction</u>. The matrix of local connectome fingerprints (841 participants × 433,386 features) contains many more features than participants (*p* ≫ *N*), thereby posing a problem for fitting virtually any type of model. To efficiently develop and evaluate predictive models in a cross-validation framework, on each fold we first performed an economical singular value decomposition (SVD) on the matrix of training subjects’ local connectome fingerprints (Wall, Andreas, and Rocha, n.d.):

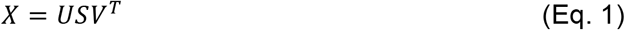

where *X* is an *n* × *p* matrix containing local connectome fingerprints for *n* participants in the cross-validation fold (~673 subjects × 433,386 elements per fingerprint), *V*^*T*^ is an *n* × *p* matrix with row vectors representing the orthogonal principal axes of *X*, and the matrix product *US* is an *n* × *n* matrix with rows corresponding to the principal components required to reproduce the original matrix *X* when multiplied by the principal axes matrix *V*^*T*^.

<u>Step 2: LASSO Model</u>. To reduce the chance of overfitting and improve the generalizability of the model for a novel test set, we employed LASSO regression, a technique that penalizes the multivariate linear model for excessive complexity (i.e., number and magnitude of nonzero coefficients) (Tibshirani, 2011). The penalty in this approach arises from the L1 sparsity constraint in the fitting process, and this combined method, known as LASSO-PCR, has been used successfully in similar high-dimensional prediction models from neuroimaging data sets (Wager et al., 2013; Wager, Atlas, Leotti, & Rilling, 2011). In short, the LASSO-PCR approach identifies a sparse set of components that reliably associate individual response variables (see Figure 1) and takes the following form:

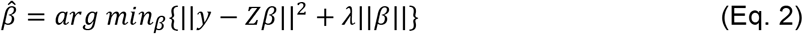

where *Z* = *US* as defined above. Using a cross-validation approach, we estimated the optimal *λ* parameter and associated 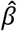 coefficients using the “glmnet” package in R (Friedman & Hastie, 2009) (see https://cran.r-project.org/web/packages/glmnet/glmnet.pdf for documentation). For each response-specific regression model, the model inputs included the principal components estimated from Eq. 1, i.e., *US* (see Figure 2), and intracranial volume (ICV). For continuous variables (e.g., reaction times), a linear regression LASSO was used. For binarized categorical variables (e.g., gender), a logistic regression variant of LASSO was used. In order to assess the value of the local connectome fingerprint components in modeling continuous response variables, the LASSO-produced 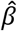 vector was truncated 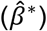 to exclude ICV and thereby restrict interpretation to the relationship between the response variables and the principal components.

The inclusion of ICV while building a model allows for the isolation of any predictive power present in the local connectome fingerprint and not to head size, which is a common adjustment used when attempting to understand structural differences between individuals or groups to reduce the possibility of type-I errors (O’Brien et al., 2011). Our LASSO-PCR procedure considers ICV in every model, and in some cases, ICV is deemed a significant contributor to variance in the response variable. In other cases, ICV is assigned a regression coefficient of zero. We observe empirically that the correlation of ICV to local connectome fingerprint principal component scores is quite small. This is to be expected considering the orthogonality of the principal components and small chance that ICV would align meaningfully with one or more component. Combining the observation that ICV has small, non-meaningful correlations with the local connectome fingerprint principal components with the knowledge that the local connectome fingerprint components are themselves orthogonal, we mitigate a common result of regression modeling in which the inclusion of a highly correlated feature may drastically alter other features’ regression coefficients. Regardless of the coefficient assigned to ICV, we ultimately want to make predictions for the continuous response variables without any knowledge of ICV by excluding the ICV coefficient and associated participant measurements from the model prediction step. While the quality of the resulting predictions (Step 4 below) may be negatively impacted by removing ICV as a potentially significant predictor in a model, controlling for ICV in this manner ensures that any observed correlation is not related to intracranial volume.

While truncating the LASSO-produced 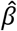 vector allows for the calculation of ICV-ignorant predictions for the continuous response variables, the same procedure cannot be adopted for categorical response variables. Such an approach to our binary responses results in undesired artifacts due to the nonlinear nature of logistic regression. An alternate approach to assess the value of the local connectome fingerprint in a binary prediction is described in Step 4.

<u>Step 3: Local Connectome Phenotype Map</u>. For each response variable, we expect 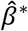 to contain non-zero weights on a subset of the orthogonal principal components (*US*, or equivalently, *XV*), and these weights were used to construct a local connectome phenotype map, defined as the weighted influence of each fiber in the local connectome on the modeled response variable. To convert the regression coefficients into the dimensions of the local connectome, the sparse vector of regression coefficients 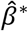 was multiplied by the principal axes matrix *V* to produce a weighted linear combination of the principal axes deemed relevant to a particular subject attribute.

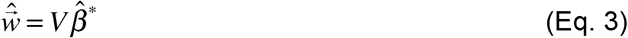

This linear combination of principal axes, 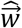, represents a *p* ×1 vector reflecting the white matter substructure of the local connectome fingerprint vector relevant to a particular observed response. We refer to the vector 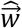 as the local connectome phenotype for the associated response variable.

<u>Step 4: Prediction</u>. Finally, we use the reconstructed local connectome phenotype map to predict a variety of continuous social, biological, and cognitive responses for participants in the test set. Ultimately, we sought a model that predicted a response variable 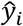 for subject *i* in the test set such that 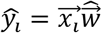 where 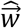 is the response-related local connectome phenotype and 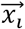 is the individual participant’s local connectome fingerprint. A prediction was generated for all participants in the hold-out set on each validation fold. Once predictions for all participants were generated for a given response variable, the performance of the model was evaluated using the correlation between predicted and observed values (continuous variables only).

While LCP maps were still constructed for categorical response variables, the utility of these LCP maps for prediction was estimated by comparing the classification accuracy of an ICV-only model to that of a model incorporating ICV and the local connectome fingerprint. In the case where the fingerprint-informed model outperforms the ICV-only model, the increase in classification accuracy can be attributed to information contained in the local connectome fingerprint map.

The calculated significance of each continuous prediction model stems from a 10,000-trial nonparametric permutation test. In each trial, the response values were permuted prior to executing the LASSO model fitting proc dure while ensuring that the fingerprint PC-ICV measurements were still paired as same-subject inputs to the models. After permuting the response values, the LASSO model fitting procedure was used to construct a response-specific model from the randomly permuted data. Correctly mapped fingerprint and ICV information was then used to predict subjects’ response values using the permutation test models. Correlation was computed for each set of model predictions and true observations to build a null distribution of the chance performance of a LASSO model for the given response. The proportion of trials in the permutation test in which the magnitude of the computed correlation met or exceeded the magnitude of the observed vs. prediction correlation in Table 3 is reported as the correlation p-value. In creating a LASSO model with permuted response values, we observed many cases in which no PCs were retained as significant predictors of variance. A resulting intercept-only model yields a constant, thus having a standard deviation of 0. Correlation between the prediction and observation in this case is undefined and was not included in the calculation of the associated p-value.

## Results

### Covariance Structure and Dimensionality of Local Connectome Fingerprints

Inter-voxel white matter architecture, reflected in the local connectome fingerprint, has been shown to be unique to an individual and sculpted by both genetic predisposition and experience (Yeh, Vettel, et al., 2016); however, it is not yet clear whether the local connectome also exhibits reliable patterns of shared variability across individuals. To illustrate this, Figure 2A shows three exemplar fingerprints from separate subjects in the sample. These exemplars reveal the sensitivity of the method to capture both common and unique patterns of variability. For example, the highest peaks in the three fingerprints are similar in terms of their size and location. This pattern appears to exist across subjects and is generally expressed in the mean fingerprint (Fig. 2C). However, there are also clear differences between participants. For example, consider the sharpness and location of the rightmost peaks in the three exemplar fingerprints in Figure 2A. This uniqueness supports our previous work highlighting single subject classification from the fingerprint across varying temporal intervals (Yeh, Vettel, et al. 2016).

**Figure 2.**
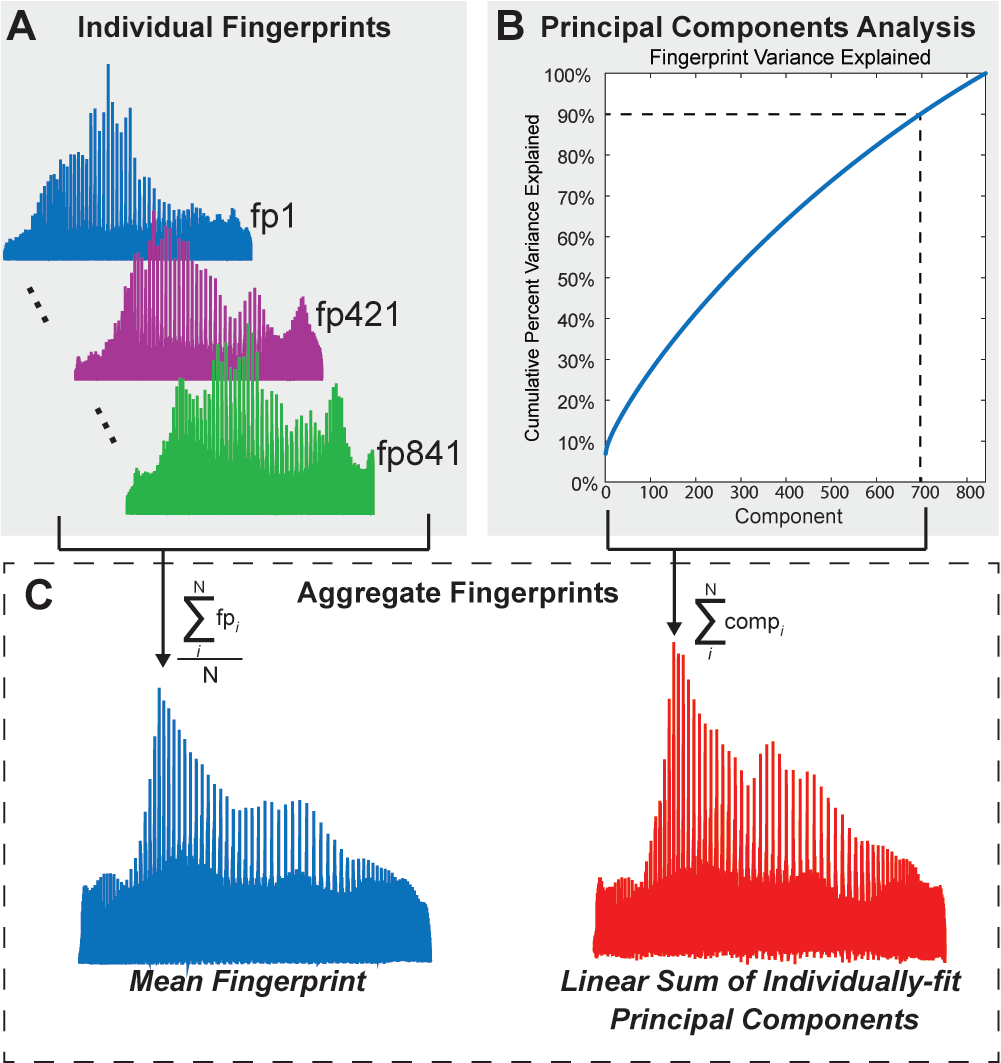
Lower dimensional structure of the local connectome fingerprints. (A) Three individual local connectome fingerprints, from three separate subjects, show coarse commonalities and unique patterns of variability when connection density is reshaped in a left-posterior-superior vector. (B) Cumulative summation of variance explained from each component, sorted by the amount of variance explained by each component. Dotted lines indicate the number of components (697) needed to explain 90% of the variability in the fingerprint dataset. (C) Mean fingerprint across participants (blue, left) and linear summation of principal components that explain 90% of the variance (red, right).

In order to explicitly test for covariance across participants, we looked at the distribution of pairwise correlations between fingerprints. The histogram in Figure 3 shows the total distribution of pairwise inter-subject correlations, revealing a tight spread of correlations such that the middle 95% of the distribution lies between 0.32 and 0.50. This confirms that intersubject correlations are substantially lower, averaging a correlation of 0.42 across all pairs of 841 HCP participants, than intra-subject correlations, found to be well above 0.90 (Yeh, Vettel, et al., 2016). Thus, the local connectome fingerprint exhibits a moderate but reliable covariance structure across participants, indicating its utility to examine shared structural variability across subjects that capture similarity in social, health, and cognitive factors.

**Figure 3.**
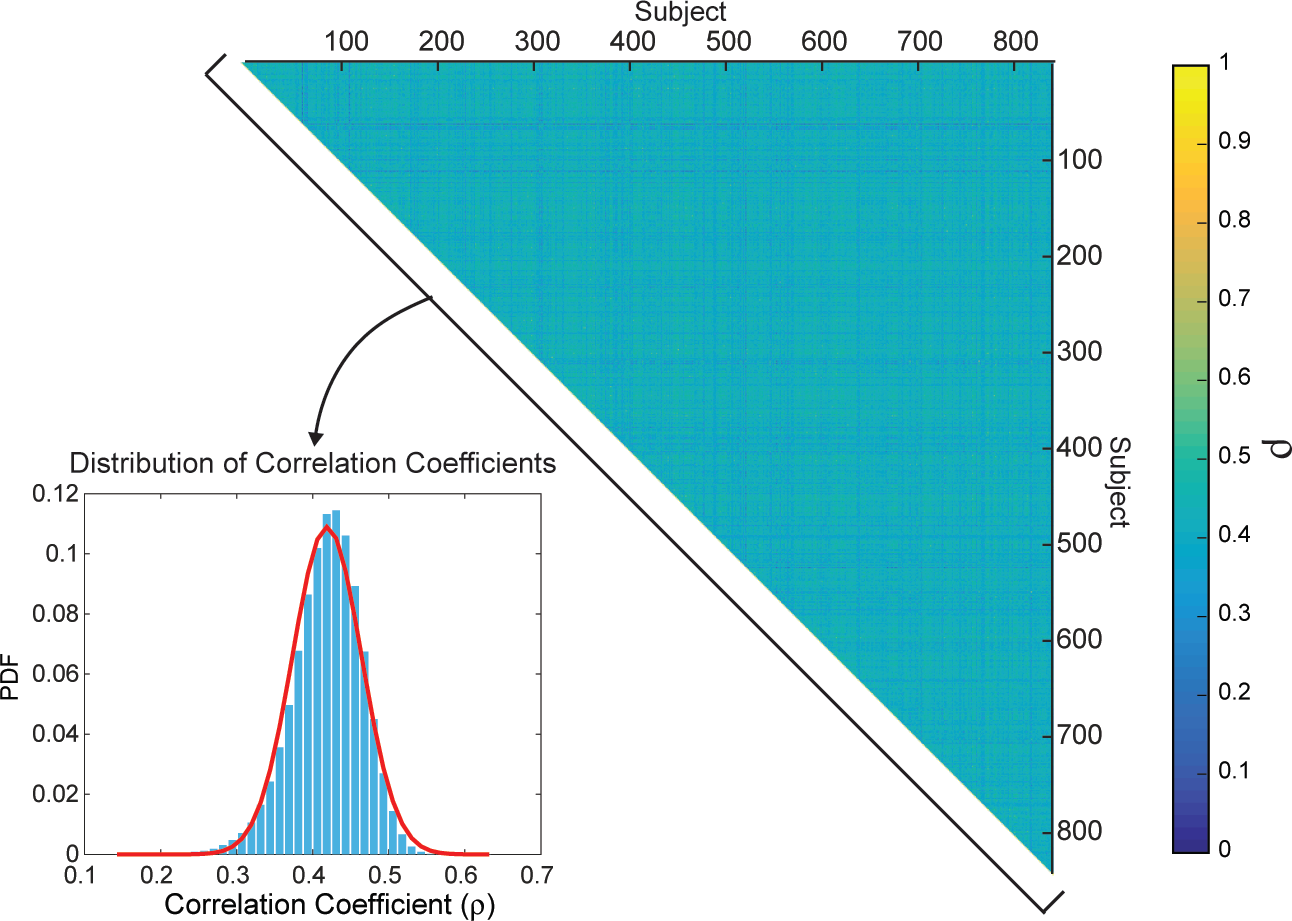
Correlations between fingerprints. The matrix of between-subject correlations in local connectome fingerprints, sorted by participant index is shown on the right. The distribution (inset) is the histogram of the upper triangle of the correlation matrix and the best fit kernel density estimate (red line).

The dimensionality of the fingerprint itself (841 participants × 433,386 elements) poses a major challenge when examining the predictive value of the local connectome for group similarity. The group fingerprint contains many more features than subjects (*p* ≫ *N*), leading to a strong risk of overfitting. We employed a dimensionality reduction routine that isolates independent principal components from the entire local connectome fingerprint matrix to decompose the variance within the set of fingerprints. This analysis found that the dimensionality of the local connectome fingerprint matrix was still relatively high and complex, requiring 697 of 841 components to explain 90% of the variance (Figure 2B). While it appears that many components are required to meaningfully explain fingerprint variance, the pattern of the mean fingerprint could be successfully recovered by a linear combination of the principal components (Figure 2C), confirming that this lower dimensional projection is adequate to represent the much larger dimensional fingerprint.

### Predicting Inter-Subject Variability

After identifying a covariance structure in the group fingerprint matrix, we fit regression models to test how well the fingerprints could predict participant attributes, including social, biological, and cognitive factors. Although we used the principal components as predictor variables, the underlying dimensionality of the local connectome fingerprint matrix (697 components for 90% variance) is still quite high relative to the sample size (841 participants). Therefore, we applied an L1 sparsity constraint (i.e., LASSO) in the fitting process of a principal components regression (LASSO-PCR), as this approach identifies a sparse set of components that reliably predict individual response variables (see Figure 1).

Table 2 shows the logistic LASSO-PCR results for the four binary categorical participant attributes: gender, race, ethnicity, and relationship status. An examination of the test accuracies in Table 2 reveals that both gender and race predictions are significantly improved with the inclusion of local connectome fingerprint information in the associated logistic regression models. The 95% confidence intervals for prediction accuracy (ICV and local connectome fingerprints) arise from bootstrapping prediction-observation pairs and reporting the appropriate percentiles from a distribution of 10,000 bootstrapped classification accuracy calculations (see Methods). The p-values associated with the reported classification accuracy arise from a nonparametric permutation test performed for each response variable. The test began by permuting response values prior to the model fitting step in order to establish a null distribution for chance accuracy achievable by a LASSO logistic regression model (see Methods). The provided p-values reflect the proportion of 10,000 trials in which the accuracy achieved in the permutation test met or exceeded the accuracy achieved in the CV-prediction of the indicated response. The models for ethnicity and relationship status revealed no relationships and perform at exactly the base rate for their respective categories.

**Table 2.** Logistic LASSO-PCR results for four categorical HCP attributes.

| Model Response (Significant CV Results *Italicized*) | Sample Size | Significant Correlation with Intracranial Volume | Training Accuracy (Measure of Model Fit) | CV-Prediction Accuracy (ICV Only) | CV-Prediction Accuracy (ICV and LCF PCs) | Confidence Interval [Lower, Upper] (ICV and LCF PCs) |  | Accuracy p-value |
| --- | --- | --- | --- | --- | --- | --- | --- | --- |
| *Gender*<sup>1*</sup> | 840 | Yes | 0.9405 | 0.8071 | 0.8691 | 0.8452 | 0.8905 | 0 |
| *Race*<sup>2*</sup> | 760 | Yes | 0.9632 | 0.8276 | 0.9053 | 0.8842 | 0.9263 | 0 |
| Ethnicity<sup>3</sup> | 833 | No | 0.9136 | 0.9136 | 0.9136 | 0.8944 | 0.9316 | 1.0000 |
| Relationship Status<sup>4</sup> | 840 | No | 0.6679 | 0.5571 | 0.5571 | 0.5226 | 0.5917 | 0.7620 |

\* The prediction accuracy was statistically significant after applying the False Discovery Rate (FDR) correction for multiple comparisons.
<sup>1</sup> The female-male split in the 840 subjects was 56%-44%, respectively.
<sup>2</sup> The white and black subpopulations made up 82% and 18%, respectively, of the 760 subjects reported here.
<sup>3</sup> The Not Hispanic/Latino and Hispanic split in the 833 subjects was 91.4%-8.6%, respectively.
<sup>4</sup> Relationship status included 44.3% of the population in a "Married or Live-in Relationship" and 55.7% not in such a relationship.

In addition to the binary participant attributes, we observed many reliable prediction models with the continuous variables. Table 3 (fourth column) shows the training results for the corresponding linear models. As expected, nearly all models were statistically significant in the training evaluation, even after adjusting for multiple comparisons. Only two variables, the Pittsburgh Sleep Quality Index and systolic blood pressure, were not significant when considering this segment of the data, largely because the LASSO model did not contain any non-zero coefficients. The LASSO form of penalized regression can drive coefficients to exactly zero when their effects are sufficiently weak. This results in an intercept-only model that produces a uniform set of predictions, and observation-prediction correlation cannot be calculated when there is no variability in the set of predictions.

**Table 3.**
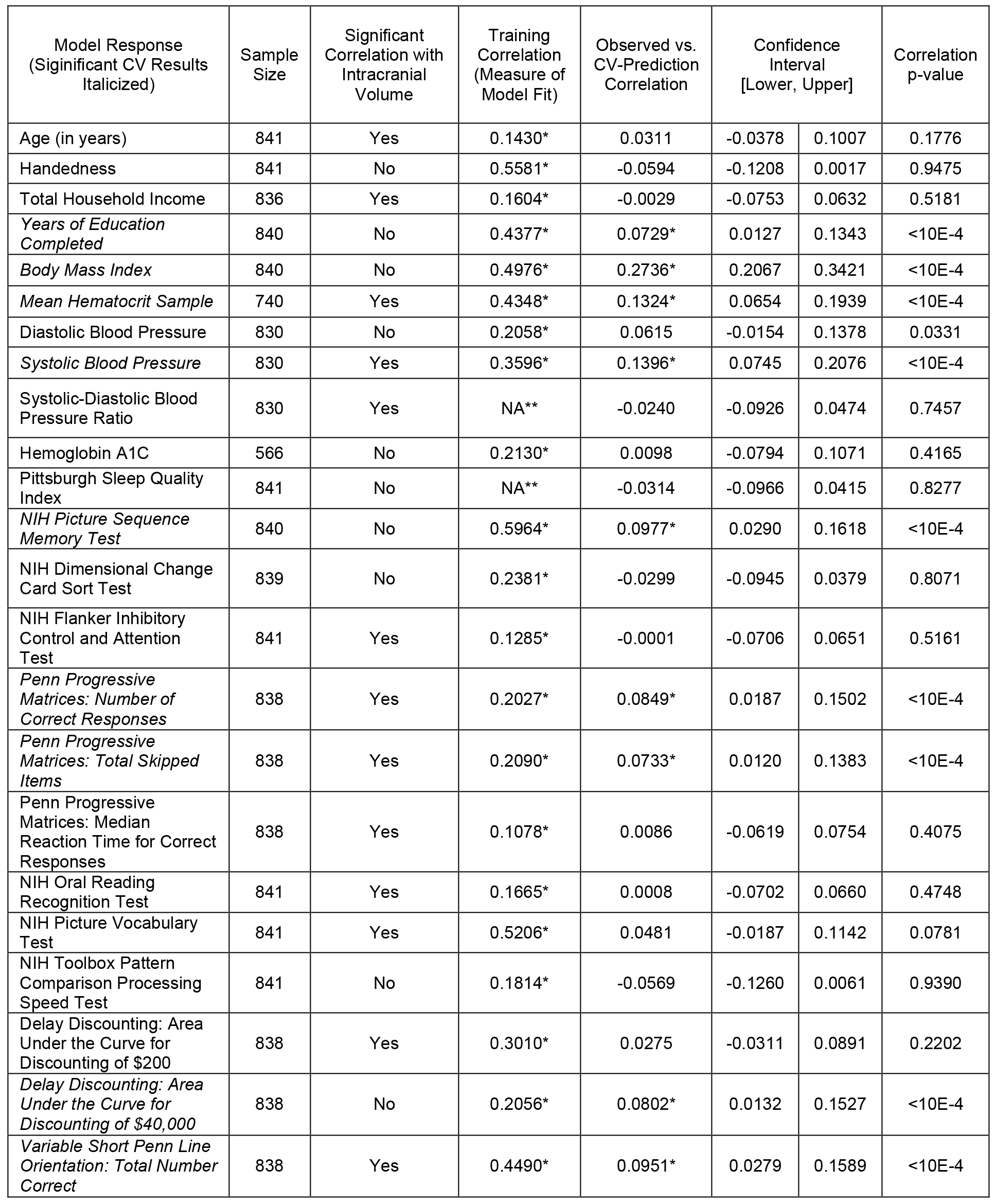
Linear LASSO-PCR results for 28 continuous HCP attributes.

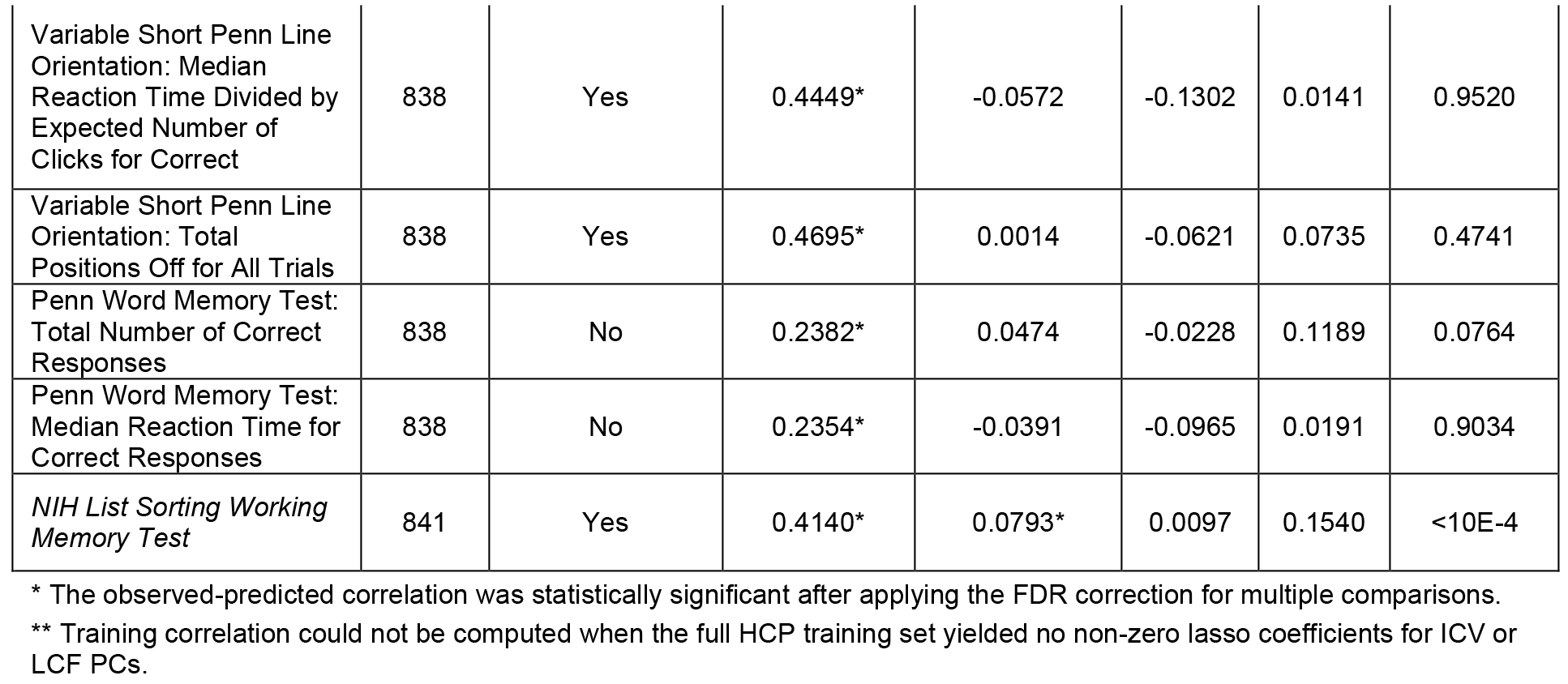

To complement the model training results, we examined the predictive performance of the models using 5-fold cross validation. This was done by projecting the regression weights in component space back into local connectome space in order to provide a weight map for each fiber in the local connectome to the target response variable. These maps reflect the local connectome phenotype for that attribute and were multiplied against a full local connectome fingerprint for each participant in the validation fold to generate a prediction for that participant (see bottom panel, Figure 1).

We assessed the generalizability of 28 continuous response models in a cross-validation paradigm and, as shown in Table 3 (fifth column), 10 of these attributes were significant predictors after applying the False Discovery Rate (FDR) correction for multiple comparisons. These factors included years of education, measures of body type (BMI), physiology (hematocrit sample, blood pressure measures), and several cognitive measures including episodic memory (NIH Picture Sequence Memory Test), fluid intelligence (Penn Progressive Matrices: Number of Correct Responses & Total Skipped Items), self-regulation (Delay Discounting: Area Under the Curve for Discounting of $40,000), spatial orientation (Variable Short Penn Line Orientation: Total Number Correct), and working memory (NIH List Sorting Working Memory Test).

### Specificity of Phenotypes to Response Variables

In our final analysis, we examined the specificity of a local connectome phenotype map by considering whether or not the predictive maps were unique for each participant attribute being predicted. In other words, we tested whether a single map could capture a generalized predictive relationship for multiple response variables, indicating that the models themselves may lack specificity. If so, any given model may perform suitably well at predicting any participant attribute (e.g., BMI), even if derived from training on a different participant factor (e.g., years of education completed).

To explicitly test this, we looked at the correlation between the 10 significant phenotype maps from the cross-validation tests shown in Table 3. This correlation is shown in Figure 4. With the exception of the correlation between the phenotypes for the Variable Short Penn Line Orientation task and the NIH List Sorting Working Memory Test, which was expected given the moderate association between performance in these two tasks (Supplementary Table 1), most of the phenotype maps were uncorrelated. We visualized the uniqueness of these phenotype maps by projecting the local connectome phenotypes into voxel space, where the average weight of multiple fibers within a voxel is depicted as a color map on the brain. A subset of these maps is shown in Figure 4. Visual inspection of these example phenotype maps reveals large heterogeneity between models. For instance, strong positive loadings are observed in portions of the splenium of the corpus callosum and frontal association fiber systems for the Picture Sequence Memory Task, while these same regions load negatively for the Variable Short Penn Line Orientation test and NIH List Sorting Working Memory Test. Bilateral corona radiata pathways appear to negatively load for the Penn Progressive Matrices and Variable Short Penn Line Orientation test, but not for any of the other attributes. These qualitative comparisons, along with the direct correlation tests, confirm that the phenotype maps for predicting intersubject variability are highly specific to the variable being modeled.

**Figure 4.**
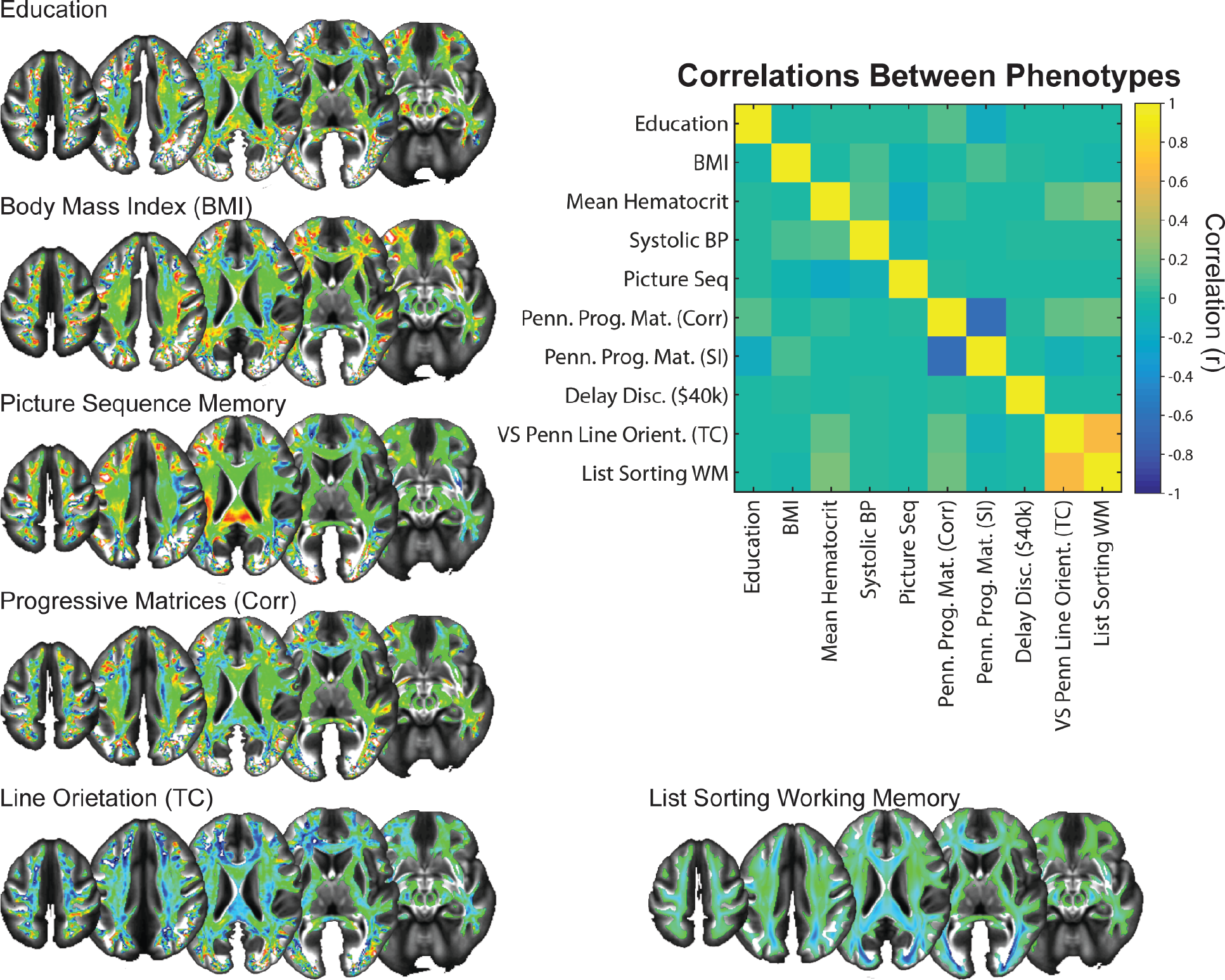
Local connectome phenotypes. Matrix inset is a correlation matrix displaying the similarity between phenotypes of the local connectome to the continuous response variables. Example phenotype maps are shown around the correlation matrix, and the color scale for each has been adjusted to reveal the areas of the local connectome that are most predictive of the labeled response variable.

## Discussion

Our analysis revealed that the local connectome fingerprint exhibits a moderate, but reliable, correlation between participants that can be leveraged to predict at the level of the individual along dimensions of social, biological, and cognitive attributes. Although the between-subject correlation is much smaller than the within-participant correlation reported previously (Yeh, Vettel, et al., 2016), it was robust enough to capture inter-subject similarities. Much to our surprise, the lower dimensional structure of this inter-subject covariance was still relatively complex, with hundreds of principal components required to explain most of the variance in the sample. Using a cross-validation regression approach that is optimized for ultra-high dimensional data sets, we show how patterns of variability in the local connectome not only correlated with nearly all participant-level social, health, and cognitive attributes (i.e., strong and significant training accuracy), but could also independently predict variability in many of the features tested (i.e., hold-out test accuracy via cross validation). Finally, we were able to show how the local connectome phenotype maps for individual attributes were highly specific to the variable being modeled. This suggests that there is not some unique, generalizable feature of local white matter that predicts inter-subject variability, but instead there are highly specific patterns that predict variance in specific inter-subject attributes. Taken together, the current results confirm our hypothesis that shared variability across participants is reflected in the local connectome itself. This opens the door for leveraging the local connectome fingerprint, along with functional measures of connectomic architecture (Shen et al., 2017), as a reliable marker for individual differences in behavior.

The current findings clearly show how it is possible to recover a portion of variability in social, biological, or cognitive attributes from local white matter architecture. This complements recent reports that global functional connectome properties can reliably predict cognitive ability (Ferguson, Anderson, & Spreng, 2017; Finn et al., 2015; Hearne, Mattingley, & Cocchi, 2016) by providing a putative structural basis for these previous associations. For example, in our study, structural similarity in the local connectome fingerprint reliably predicted six of the tested cognitive performance measurements, including a list sorting task that captures individual variability in working memory performance (Gur et al., 2001; Gur et al., 2010). The associated local connectome phenotype for working memory identified portions of what appear to be frontoparietal pathways (Figure 4). Our results nicely complement a recent study of working memory that focused on direct and indirect connectivity in the frontoparietal networks (Ekman, Fiebach, Melzer, Tittgemeyer, & Derrfuss, 2016). In their work, the authors found that the network centrality of focal structural connections in the frontal, temporal, and parietal cortices could predict individual differences in working memory capacity using linear regression. When considered in the context of the current study, our findings augment previous correlative findings between frontoparietal regions and working memory capacity (Bender, Prindle, Brandmaier, & Raz, 2016; Klingberg, 2006; Nagy, Westerberg, & Klingberg, 2004; Takeuchi et al., 2010) by showing that the integrity of the pathway of these white matter fascicles reliably predicts working memory performance.

The existence of reliable and predictive inter-subject covariance patterns in the white matter fascicles of the human brain begs the question of mechanism: are these similarities genetically determined, experientially sculpted, or developed through gene-by-environment interactions? Emergent findings in genetics are suggesting that at least a portion of macroscopic white matter structure is guided by genetics (Kochunov, Fu, et al., 2016; Kochunov, Thompson, et al., 2016; Yeh, Vettel, et al., 2016). For example, recent work by Kochunov and colleagues (2016a) examined a heritability relationship between whole-brain fractional anisotropy (FA) and information processing speed in two interesting participant populations, the HCP twins cohort and an Old Order Amish cohort. The cohorts both had well-characterized genetic properties, but they differed in the amount of experiential variability since the Amish have higher environmental homogeneity compared to the urban/suburban HCP cohort. Kochunov and colleagues (2016a) argued that the replication of the genetic contribution to processing speed and FA of cerebral white matter despite the experiential variability in the cohorts suggested a strong phenotypic association for the trait. Our analysis would be able to pick up such genetically mediated brain-behavior phenotypes.

While genetics may contribute to white matter architecture, overwhelming evidence suggests that experience sculpts these pathways over time. For example, variability in the white matter signal has been shown to covary with several social (Gianaros et al., 2013; Molesworth et al., 2015), biological (Arfanakis et al., 2013; Miralbell et al., 2012; Verstynen et al., 2013), and cognitive (Muraskin et al., 2016; Verstynen, 2014; Ystad et al., 2011) attributes. In many cases, it is difficult to extract or identify specific pathways or systems that link white matter pathways to these shared experiential factors. However, several intervention studies have targeted more specific experience-white matter associations. For example, prolonged training on a variety of tasks has been shown to induce changes in the diffusion MRI signal (Blumenfeld-Katzir, Pasternak, Dagan, & Assaf, 2011; Sampaio-Baptista et al., 2013; Scholz, Klein, Behrens, & Johansen-Berg, 2009; Steele, Scholz, Douaud, Johansen-Berg, & Penhune, 2012). In some cases, the particular change in the diffusion signal is consistent with alterations in the underlying myelin (Sampaio-Baptista et al., 2013), for which there is emerging support from validation studies in non-human animal models (Budde, Janes, Gold, Turtzo, & Frank, 2011; Budde, Xie, Cross, & Song, 2009; Klawiter et al., 2011). One consistency in these reports of training-induced plasticity in white matter pathways is that the effects are task-specific (i.e., training in a specific task appears to impact specific white matter fascicles). This specificity of experiential factors on white matter pathways is necessary in order to be able to build reliable prediction models from the diffusion MRI signal.

Our previous work showed that the local connectome fingerprint reflects both genetic and experiential factors that contribute to between-subject variability in white matter architecture (Yeh, Vettel, et al., 2016). We found that monozygotic twins expressed a modest degree of similarity in their local connectome fingerprints, with ~12% of the local connectome pattern being similar between monozygotic twins. This similarity was much higher than what was detected in siblings or dizygotic twins; however, genetic similarities overall seemed to contribute very little to similarities in the local connectome. In contrast, most of the structure in the local connectome fingerprint appeared to be driven by experience. By comparing changes in the fingerprint over time, average intra-subject similarity changed linearly with time. While it can be argued that part of this change simply reflects aspects of the normal aging process (Simmonds, Hallquist, Asato, & Luna, 2014; Westlye et al., 2010), we should point out that the intra-subject changes seen in our previous study happen at a much faster rate than typical age-related changes in white matter pathways (i.e., days and weeks vs. years, respectively). Thus, we expect that much of this plasticity is likely due to experiential factors.

One of the strengths of the local connectome fingerprint approach used here is that it does not rely on fiber tracking algorithms. Recent evidence indicates a false positive bias when mapping white matter pathways (Daducci et al., 2016; Reveley et al., 2015; Thomas et al., 2014). This is due in large part to the difficulty that tracking algorithms have when distinguishing between a crossing and turning fiber pathway. Our approach does not rely on a deterministic or probabilistic tracking algorithm; instead, we analyze the entire set of reconstructed fibers throughout the brain as a unitary data object. This eliminates the false positive identification of white matter fascicles by not attempting fascicular classification at all. However, without tracking along pathways we cannot say whether specific pathways positively or negatively predict a specific response variable. In the future, exploration of the local connectome phenotype maps with careful pathway labeling (e.g., expert-vetted fiber labeling) can identify general regions that positively or negatively contribute to the prediction.

Another limitation of the approach used here arises from the fact that, by necessity, the local connectome fingerprints must be computed from a common, atlas-defined space. The nonlinear transformations required in order to transform brains of various shapes and sizes into a stereotaxic space through the QSDR procedure invariably introduce a degree of noise in the SDFs. The number and orientation of fibers in each voxel determine the local connectome fingerprint, and these measurements could possibly be distorted during QSDR. Such a transformation is unavoidable because the dimensionality of each fingerprint must be identical, and each element of a fingerprint must represent the same brain micro-region as the corresponding element in any other fingerprint. Only with this common, atlas-aligned representation of the local connectome fingerprint can we apply LASSO-PCR to explore common substructures. The potential price for this convenience is an introduction of noise in the local connectome fingerprint itself, likely increasing the possibility of a false-negative error (i.e., failing to recognize a true phenotypic relationship). In addition, the sampling of the local connectome comes from identifying the peaks from the average SDF for this particular sample of healthy young adults. While it is believed that this approach gives a reasonable estimate of normative fiber structure (Yeh, Vettel, et al., 2016), it is possible that an atlas defined from another population, with consistent differences in local white matter architecture (e.g., older adults), could result in slightly different local connectome fingerprints and thus slightly different phenotypic associations.

Our analytical design was constructed to examine the generalizability of associations between local white matter architecture and demographic, health, and cognitive attributes rather than to investigate simple descriptive correlations. Although training accuracies themselves do not evaluate how well the model generalizes to unseen data, we included training model performance results in Tables 2 and 3 to highlight two important points. First, in some cases, test model performance is poor because the training model is also poor. This reflects cases where the model fitting procedure simply failed to identify meaningful patterns, as opposed to cases where the fitting procedure was highly biased to the training set, but exhibits low flexibility (i.e., sensitive to meaningful, but not generalizable associations). Second, and more importantly, many traditional neuroimaging approaches only report training model results that often overestimate the strength of the relationship. Results in Tables 2 and 3 reveal that nearly all training models show strong, significant associations; however, only a small subset retain significance on the independent hold out set, where the effect size is much smaller. We should note that the effect sizes of the significant models in the hold out test validation, particularly the cognitive measures, are substantially smaller than previously reported effect sizes of functional connectome phenotypes (Ferguson, Anderson, & Spreng, 2017; Finn et al., 2015; Hearne, Mattingley, & Cocchi, 2016). This may be due to the fact that variability in structural connections may serve as a moderator of global network dynamics that drive behavior, but the functional dynamics themselves are a more direct reflection of immediate brain function. This suggests that multimodal analysis accounting for both structural and functional connectomic architecture may provide a stronger prediction of individual variability in cognitive function.

The current work reveals that the local connectome fingerprint reliably reflects shared variance between individuals in the macroscopic white matter pathways of the brain. For the first time, we not only show how global white matter structure associates with different participant features, but we also show how the entire local connectome itself can predict a portion of the variability in independent samples. While the overall variance explained by the local connectome fingerprint may at first seem small, it is consistent or even stronger than effect sizes of genetic risk scores used in behavioral medicine (Plomin, DeFries, Knopik, & Neiderhiser, 2016). Thus, our local connectome phenotyping approach may also be predictive of not only normal, but also pathological variability (see also Yeh et al., 2013). Future work in clinical populations should focus on applying this approach to generate diagnostic local connectome phenotypes for neurological and psychiatric disorders, thereby leveraging the full potential of this approach.

---

**Supplementary Table 1.**
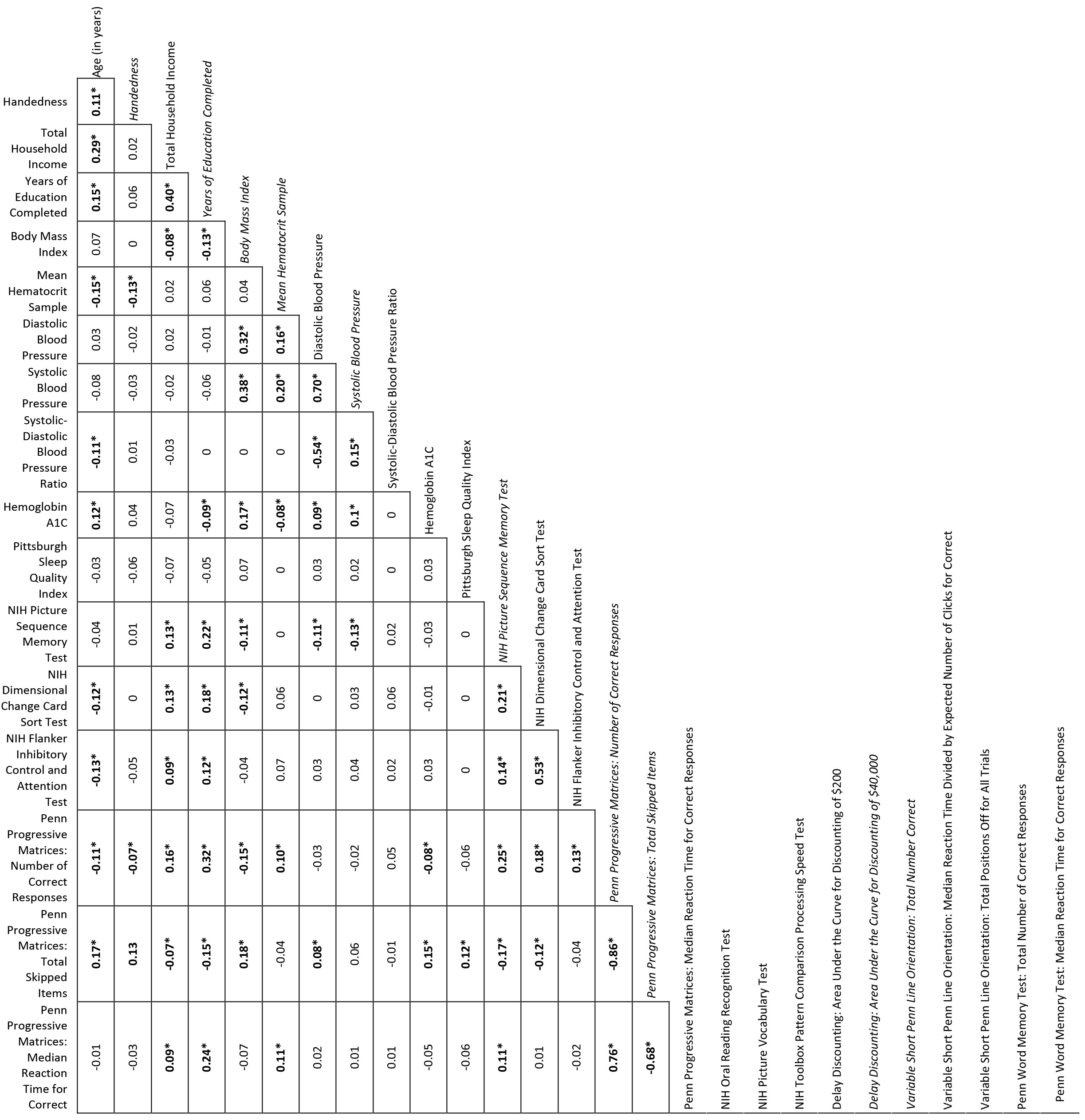
Pairwise Pearson correlation matrix for 28 continuous HCP attributes. Asterisks indicate significant correlations after correcting for multiple comparisons (FDR < 0.05).

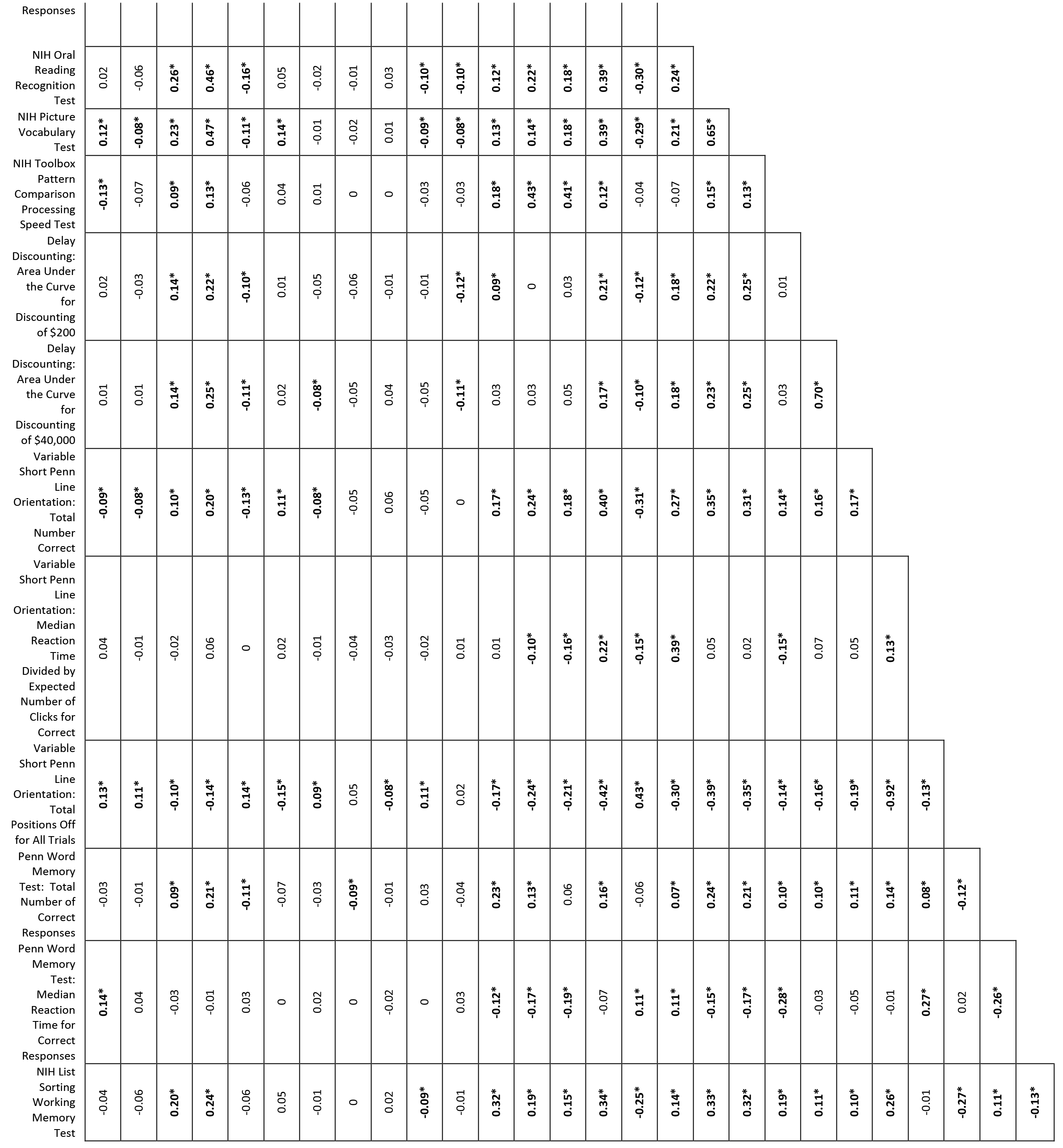

## Acknowledgements

The research was sponsored by the Army Research Laboratory and accomplished under Cooperative Agreement Number W911NF-10-2-0022. The views and conclusions contained in this document are those of the authors and should not be interpreted as representing the official policies, either expressed or implied, of the Army Research Laboratory or the U.S. Government.

